# Read trimming has minimal effect on bacterial SNP calling accuracy

**DOI:** 10.1101/2020.08.04.236216

**Authors:** Stephen J. Bush

## Abstract

Read alignment is the central step of many analytic pipelines that perform SNP calling. To reduce error, it is common practice to pre-process raw sequencing reads to remove low-quality bases and residual adapter contamination, a procedure collectively known as ‘trimming’. Trimming is widely assumed to increase the accuracy of SNP calling although there are relatively few systematic evaluations of its effects and no clear consensus on its efficacy. As sequencing datasets increase both in number and size, it is worthwhile reappraising computational operations of ambiguous benefit, particularly when the scope of many analyses now routinely incorporate thousands of samples, increasing the time and cost required.

Using a curated set of 17 Gram-negative bacterial genomes, this study evaluated the impact of four read trimming utilities (Atropos, fastp, Trim Galore, and Trimmomatic), each used with a range of stringencies, on the accuracy and completeness of three bacterial SNP calling pipelines. We found that read trimming made only small, and statistically insignificant, increases in SNP calling accuracy even when using the highest-performing pre-processor, fastp.

To extend these findings, we re-analysed > 6500 publicly-archived sequencing datasets from *E. coli*, *M. tuberculosis* and *S. aureus*. Of the approximately 125 million SNPs called across all samples, the same bases were called in 98.8% of cases, irrespective of whether raw reads or trimmed reads were used. However, when using trimmed reads, the proportion of non-homozygous calls (a proxy of false positives) was significantly reduced by approximately 1%. This suggests that trimming rarely alters the set of variant bases called but can affect their level of support. We conclude that read quality- and adapter-trimming add relatively little value to a SNP calling pipeline and may only be necessary if small differences in the absolute number of SNP calls are critical. Read trimming remains routinely performed prior to SNP calling likely out of concern that to do otherwise would substantially increase the number of false positive calls. While historically this may have been the case, our data suggests this concern is now unfounded.

**Impact Statement:** Short-read sequencing data is routinely pre-processed before use, to trim off low-quality regions and remove contaminating sequences introduced during its preparation. This cleaning procedure – ‘read trimming’ – is widely assumed to increase the accuracy of any later analyses, although there are relatively few systematic evaluations of trimming strategies and no clear consensus on their efficacy. We used real sequencing data from 17 bacterial genomes to show that several commonly-used read trimming tools, used across a range of stringencies, had only a minimal, statistically insignificant, effect on later SNP calling. To extend these results, we re-analysed > 6500 publicly-archived sequencing datasets, calling SNPs both with and without any read trimming. We found that of the approximately 125 million SNPs within this dataset, 98.8% were identically called irrespective of whether raw reads or trimmed reads were used. Taken together, these results question the necessity of read trimming as a routine pre-processing operation.

**Data Summary:** All analyses conducted in this study use publicly-available third-party software. All data and parameters necessary to replicate these analyses are provided within the article or through supplementary data files. > 6500 SRA sample accessions, representing Illumina paired-end sequencing data from *E. coli*, *M. tuberculosis* and *S.aureus*, and used to evaluate the impact of fastq pre-processing, are listed in **Supplementary Tables 3**, **5 and 7**.

## Introduction

Read alignment is the central step of many analytic pipelines that perform variant calling. To reduce error, it is common practice to pre-process raw sequencing reads to remove low-quality bases and residual adapter contamination, a procedure collectively known as ‘trimming’. This is because, assuming Illumina sequencing data, errors are non-randomly distributed over the length of the read, clustering towards the 3’ end (which is also where adapters are located). These poorer-quality flanking regions are frequently trimmed to leave only the higher-quality internal bases.

Numerous pre-processing tools (“read trimmers”) exist for this purpose, which often simultaneously perform both quality- and adapter-trimming. However, previous studies differ as to whether the effect of trimming on downstream SNP calling is generally beneficial (1, 2), generally minimal (3), or conditional on the genome and aligner used (4). Similarly, trimming was reported to have little effect on the completeness of *de novo* genome assembly (5) and be detrimental to *de novo* transcriptome assembly unless comparatively gentle (i.e. trimming on the basis of quality score < 5) (6). Taken together, the benefits of trimming do not appear universal and in many situations may not be realised at all.

As sequencing datasets increase both in number and size, it is worthwhile reappraising computational operations of ambiguous benefit, particularly when the scope of many analyses now routinely incorporate thousands of samples. To that end, this study evaluates the effect of several read trimming strategies on the subsequent accuracy and completeness of various bacterial SNP calling pipelines. We evaluated four commonly-used read trimmers – fastp (7), Trimmomatic (8), Atropos (9), and TrimGalore (the latter two employing Cutadapt (10)), each of which applies a different strategy to adapter detection and removal – across a range of stringencies.

To assess the effect of pre-processing stringency upon the precision (positive predictive value) and recall (sensitivity) of SNP calling, we quality- and adapter-trimmed 17 sets of 150bp Illumina HiSeq 4000 paired-end reads before calling SNPs relative to a set reference genome using three different pipelines (the pairwise combination of one read aligner, BWA-mem (11), and three variant callers, LoFreq (12), mpileup (13), and Strelka (14)). These reads represent environmentally-sourced samples of the genera *Citrobacter*, *Klebsiella*, *Escherichia*, and *Enterobacter* and were obtained from a previous study (15) and curated for use in a large-scale comparison of bacterial SNP calling pipelines (16). The three pipelines chosen for the present study were previously found to be among the highest-performing when tested on divergent bacterial data (16).

To determine the effect of trimming on a larger-scale dataset, we then applied the highest performing strategy identified for the curated Gram-negative dataset to a set of > 6500 publicly archived *Escherichia coli*, *Mycobacterium tuberculosis* and *Staphylococcus aureus* sequencing reads. This represents a substantive, and diverse, range of Illumina sequencing platforms, library preparation strategies, read lengths, insert sizes, and coverage depths. In this dataset, we have no *a priori* knowledge of which SNP calls are accurate, although for the purpose of this analysis this is not relevant – the intention was to compare two sets of SNPs called before and after a common set of pre-processing steps were applied, and so we were not interested in the accuracy of each call but whether the same calls were made in each condition. This dataset is sufficiently large (approximately 125 million SNPs) that we can generalise about the benefits of read trimming as a routine procedure.

## Results and Discussion

### Read trimming has minimal effect on SNP calling accuracy

We have previously described how the performance of a bacterial SNP calling pipeline is affected by divergence between the genome from which reads are sequenced and the genome to which reads are aligned, the latter often being the NCBI ‘reference genome’, a high-quality (albeit often arbitrary) species representative (16). To do so, we obtained data from a diverse range of environmentally-sourced Gram-negative bacteria (as described in (15) and summarised in **Supplementary Table 1**) and used it to generate truth sets of SNPs for evaluation purposes, as previously detailed (16) (and briefly recapitulated in the Materials and Methods). We chose three SNP calling pipelines found to be generally higher-performing, even when reads were aligned to particularly divergent genomes (16): the pairwise combination of the aligner BWA-mem (11) with three different variant callers, LoFreq (12), mpileup (13), and Strelka (14), each used with a common set of post-alignment processing operations.

We used these three pipelines to call SNPs across the range of Gram-negative bacteria, both in the absence of read pre-processing and after preprending to each pipeline one of four different read trimmers: Atropos, fastp, TrimGalore and Trimmomatic. Each trimmer was configured to remove adapter sequence as well as to trim bases from the 3’ end of each read, on the basis of Phred quality score (Q) thresholds 2, 5, 10, 20 and 30 (the Phred scale being logarithmic, this represents base call accuracies of 37%, 68%, 90%, 99%, and 99.9%, respectively). In total, this dataset contained 1071 records, comprising 17 species x 4 read trimmers x 5 Phred score thresholds x 3 variant callers, plus 17 species x 3 variant callers provided untrimmed data (i.e. not including the latter, 50 records per trimmer/Phred threshold combination).

We found that across the full range of genomes and aligner/caller combinations, trimming had minimal effect on overall SNP calling accuracy, with only miniscule, insignificant, changes observed in F-score relative to untrimmed data (these changes visible only in the third decimal place of F-score; see **Figure 1** and **Supplementary Table 2**). It appeared that the highest-performing trimming strategy employed fastp which, relative to untrimmed data, produced consistent, albeit statistically insignificant, increases in F-score irrespective of trailing Phred threshold (the median F-score using untrimmed data was 0.9579 and, using fastp with a threshold of Q < 20, 0.958; Mann-Whitney U p = 0.976). This pattern appears driven by relatively small decreases in precision (**Supplementary Figure 1**) compensated by slightly larger increases in recall (**Supplementary Figure 2**), which suggests that fastq pre-processing enables a small proportion of reads to map that otherwise would not, allowing lower-coverage SNPs to be called that would otherwise be omitted. It is important to note that fastp differs from the other trimmers in that, unless explicitly disabled, it implements several filters by default – that is, it performs more quality-trimming operations than simply 3’ trimming (discussed below). Notably, 3’ trimming alone (as implemented by the other three trimmers) appears to uniformly, albeit marginally, improve precision (**Supplementary Figure 1**) although at the expense of recall (**Supplementary Figure 2**) and thereby overall F-score (**Figure 1**). This is also the case for fastp if repeating the analysis with the default filters disabled (**Supplementary Figure 3**).

**Figure 1.**
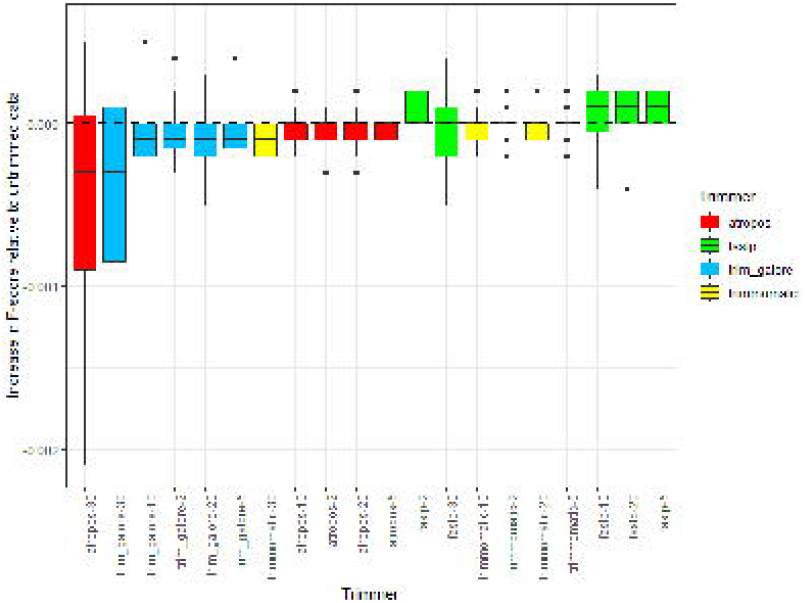
Effect of read trimming upon F-score, a measure of overall SNP calling accuracy, in a curated Gram-negative dataset. Median difference in F-score per trimmer relative to untrimmed data, across a range of trimming stringencies (i.e. varying the Phred score threshold for trimming 3’ bases). Boxes represent the interquartile range of F-score, with midlines representing the median. Upper and lower whiskers extend, respectively, to the largest and smallest values no further than 1.5x the interquartile range. Data beyond the ends of each whisker are outliers and plotted individually. Columns are ordered according to median F-score and coloured according to the trimmer used. The dashed line y=0 is marked in black. The raw data for this figure is available in **Supplementary Table 2**. Boxplots showing the effect of read trimming upon precision and recall are shown, respectively, in **Supplementary Figures 1 and 2**. Note that fastp implements quality filters other than 3’ trimming by default, which for the data in this figure were retained. A version of this figure with these filters disabled is available in **Supplementary Figure 3**.

Nevertheless, if considering SNP calling in absolute terms, these results suggest that there is no overt disadvantage to trimming, but little substantive benefit either. To ensure this conclusion is not generalised on the basis of a small number of samples, we selected one SNP calling pipeline (BWA-mem/mpileup) and then applied one of the higher-performing trimming strategies (fastp with trailing Q < 20) to hundreds of publicly sourced, and genomically diverse, *E. coli* samples (**Supplementary Table 3**), calling SNPs in all cases relative to a single reference genome and applying a common set of post-processing operations to restrict analysis only to higher-confidence calls (see Materials and Methods). We found that trimming, in general, improved the proportion of reads that could be aligned (**Figure 2A**) although SNP calling – that is, the interpretation of those alignments – was not substantially altered. In 1579 of the 1606 *E. coli* samples, > 99% of SNPs could be identically called irrespective of read trimming (> 99.9% in 385 samples, 24% of the total) (**Figure 2B** and **Supplementary Table 4**). Of the total set of > 103 million *E. coli* SNPs, 99.85% were identically called irrespective of any read trimming. It is important to clarify that by ‘identical’ we are referring only to whether the base calls are the same, e.g. G/A, and do not distinguish here between a G/A homozygote and a G/A heterozygote, this being a matter of VCF post-processing (see below).

**Figure 2.**
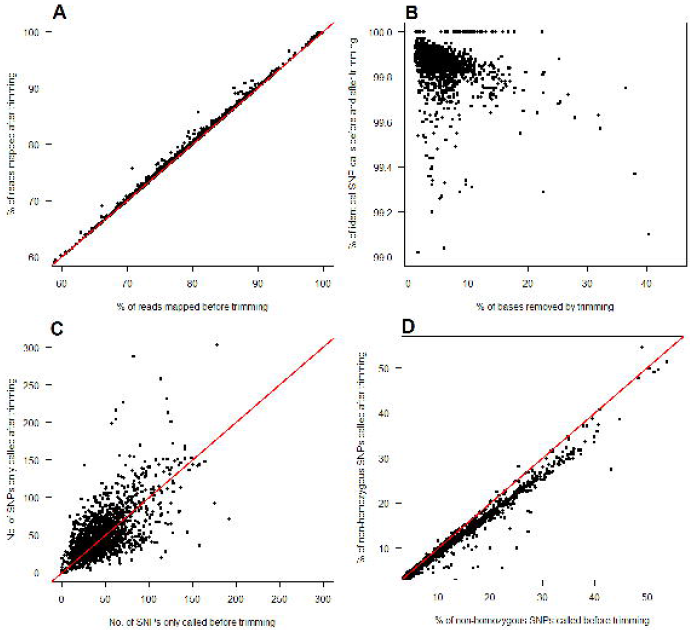
Effect of read trimming upon SNP calls made using publicly-archived *E. coli* sequencing data. Trimming marginally increases the proportion of successfully aligned reads (panel A) although the interpretation of those alignments (i.e. SNP calling) is not substantially altered, with the vast majority of SNPs (> 99%) called irrespective of trimming (panel B). This value is 100% for a number of samples containing very few SNPs (approximately 15) relative to the *E. coli* reference genome. A relatively small number of SNPs (in the majority of cases, < 200) are only called when using either raw, or trimmed data, but not both (panel C). The proportion of non-homozygous SNP calls, considered a proxy of false positive calling, decreases when using trimmed data (panel D). The raw data for this figure is available in **Supplementary Table 4** and represents 1606 *E. coli* samples, with a mean of 64,476 SNPs per sample. The red line denotes y = x.

The negative correlation between the percentage of bases removed by trimming and the percentage of identically called SNPs (**Figure 2B**) likely reflects reduced sequencing depth in the more extreme cases – some samples are evidently lower-quality (according to the filter criteria applied), with > 20% of bases trimmed. In one of the more extreme cases, sample SRS3938880 (collected during a transnational survey of veterinary pathogens (17)), 40% of its original set of 1000 million bases were discarded, although compared to the raw data, 99.1% of the total set of 53,701 SNPs could still be called (**Supplementary Table 4**). In absolute terms it also appears that approximately as many SNPs per sample were called only when using trimmed reads as when using raw reads (**Figure 2C**). However, even if we were to assume that every SNP called only when using raw reads was erroneous (that is, it is an error that trimming will resolve), this remained < 200 SNPs for 1598 of 1606 samples (**Supplementary Table 4**).

Although we were unable to ascertain the accuracy of any of the SNP calls in the publicly-sourced dataset, and thereby which were true positives, we can nevertheless assume that non-homozygous calls (those where < 100% of reads mapped at that position support the variant allele) are a proxy of false positives, as in a previous study (1). As the proportion of non-homozygous SNP calls is marginally higher for raw than trimmed data, this suggests that SNP calling using raw data may introduce a number of errors that trimming would resolve (the median percentage of non-homozygous SNP calls is 10.26% when using untrimmed data, and 9.05% when using trimmed data; Mann-Whitney U p = 9.1×10^−7^; **Figure 2D**). Taking the above results together, we can conclude that compared to raw reads, the use of trimmed reads rarely changes the variant base called (see **Figure 2B**) but does alter the level of support for a given variant in approximately 1% of cases. However, it is worth noting that the identification of (potentially spurious) non-homozygous calls is by definition a VCF post-processing operation and so could be applied regardless of any fastq pre-processing, and at varying levels of stringency.

*E. coli* is a characteristically diverse species, and so in this analysis the mean number of SNPs per sample was relatively high, at approx. 70,000 (**Supplementary Table 4**). A consequence of this genomic diversity is that it minimises the variance introduced by small differences in SNP calling, before and after trimming, which in another situation could be critical. To explore the effect of trimming on a clonal system, where only a small number of SNPs are expected (and so trimming-associated differences would have greater impact), we repeated the analysis using 3946 publicly-sourced *M. tuberculosis* samples (**Supplementary Tables 5 and 6**).

Unlike *E. coli*, which is sufficiently diverse that only approximately 75% of reads per sample could be aligned to the reference genome (this proportion marginally increased after trimming; **Figure 2A**), virtually all *M. tuberculosis* reads could be successfully aligned to the reference, H37Rv, regardless of trimming (although as with *E. coli*, trimming also increased the proportion of aligned reads, as shown in **Figure 3A**). Similar results were observed for *M. tuberculosis* as for *E. coli* with, in the majority of cases, > 98% of SNPs identically called irrespective of read trimming (**Figure 3B**), a small number of SNPs (often < 50) only called when using raw reads or trimmed reads, but not both (**Figure 3C**), and a small but significant decrease in the number of non-homozygous calls made after trimming (the median percentage of non-homozygous calls is 18.9% when using untrimmed data, and 14.1% when using trimmed data; Mann-Whitney U p < 2.2×10^−16^; **Figure 3D**). Quantitatively similar results were again observed if expanding the analysis to incorporate 1100 publicly-archived *S. aureus* samples (**Supplementary Tables 7 and 8**, and **Supplementary Figure 4**), with the exception that there was no significant decrease in the number of non-homozygous calls made by trimming (the median percentage of non-homozygous calls is 12.7% when using untrimmed data, and 12.6% when using trimmed data; Mann-Whitney U p = 0.48; **Supplementary Figure 4**). Across the combined set of approximately 125 million SNPs from all three species, 98.8% of SNPs could be identically called irrespective of any read trimming.

**Figure 3.**
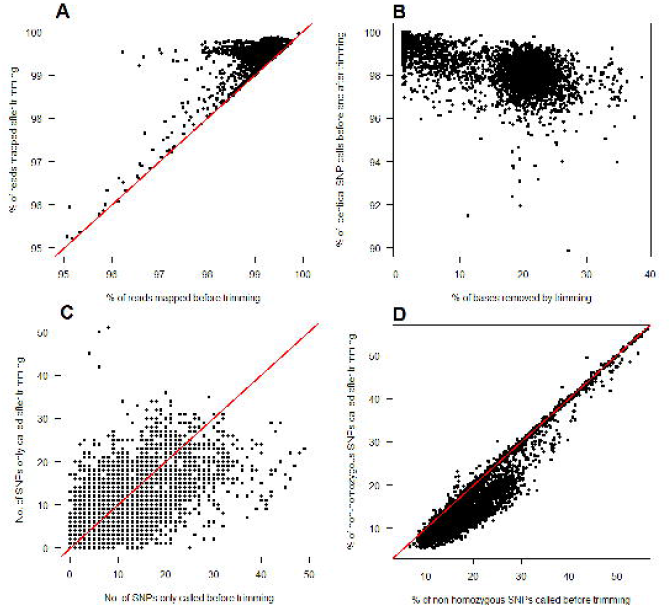
Effect of read trimming upon SNP calls made using publicly-archived *M. tuberculosis* sequencing data. This figure recapitulates patterns seen in **Figure 2** and illustrates the effect of read trimming upon SNP calls made in a clonal species, *M. tuberculosis*, for which relatively high alignment accuracy is expected and the impact of misalignment (i.e. false positive SNP calls) accordingly greater. Trimming marginally increases the proportion of successfully aligned reads, albeit from a high baseline value, > 98% (panel A). The vast majority of SNPs (> 98%) are nevertheless called irrespective of any trimming (panel B). A relatively small number of SNPs (often < 40) are only called when using either raw, or trimmed data, but not both (panel C). The proportion of non-homozygous SNP calls, considered a proxy of false positive calling, decreases when using trimmed data (panel D). The raw data for this figure is available in **Supplementary Table 6** and represents 3946 *M. tuberculosis* samples, with a mean of 1238 SNPs per sample. The red line denotes y = x.

### Recommendations for read trimming prior to SNP calling

We found that read trimming, in general, had minimal effect on the performance of a BWA-mem/mpileup SNP calling pipeline. This suggests that routine read trimming is not always a practical necessity. If the purpose of SNP calling is to construct bacterial phylogenies (for instance, to infer transmission) then trimming appears of little value given the majority of SNPs in a sample are identically called regardless of whether reads are trimmed or not. By contrast, if reducing the likelihood of even a small number of false positive calls is essential (for instance, when predicting antimicrobial resistance) then trimming may yet prove critical. Supporting this point, we found that after re-analysing a large number of publicly archived datasets there were small (c. 1%) but significant decreases in the proportion of non-homozygous SNP calls made when using trimmed compared to untrimmed data (**Figures 2D and 3D**), these considered a proxy for false positive calls.

Many of the justifications for routine read trimming relate to its simplicity: it is an easily performed procedure. In this respect, if it is only necessary to process a limited number of samples and computational resources are not at a premium, there appears little detriment to trimming – albeit, as we have shown, not necessarily much benefit either.

However, if computational resources are at a premium, then various factors may be taken into consideration before attempting to trim reads. Firstly, when a trimmer is provided paired-end reads as input, it can produce both paired- and single-end reads as output, the latter when only one end of a pair is discarded. Directing paired and unpaired reads to separate sets of output files is necessary as in general a read aligner, if aligning paired-end reads, requires an identical number of reads in each input fastq. However, as numerous aligners do not allow the simultaneous provision of paired-end and unpaired input (with some exceptions, such as HISAT2 (18)), any subsequent alignment step would need to be run twice and the output BAMs merged, an additional (and therefore time-consuming) set of operations. The default parameters of fastp and TrimGalore are to output only the trimmed paired-end reads which, although convenient, does discard a small proportion of otherwise useable single-end data. We found, however, that these single-end reads were often few in number and negligibly informative and so in many cases could reasonably be dispensed with. We re-processed the 1606 *E. coli* samples, discarding those reads unpaired after trimming, and found negligible difference in the total number of SNPs called using only paired-end reads and using all reads (the median number of SNPs called when only using paired-end reads and when using all reads are 55,137 and 55,147, respectively; Mann-Whitney U p = 0.871; **Supplementary Figure 5**). There was also a small decrease in the number of non-homozygous calls made when discarding the unpaired reads, although this difference was not significant (the median percentages of non-homozygous calls made when retaining, and discarding, unpaired reads were 9.03% and 8.51%, respectively; Mann-Whitney U p = 0.055) (**Supplementary Figure 5**).

Secondly, we need to consider the computational cost of pre-processing. If calling SNPs for a large number of samples, this cost may be weighed negatively against the relative speed and simplicity of post-processing a VCF – for instance, by masking repetitive regions and applying positional filters (both quick operations), SNP calling precision can easily be increased (19). We reasoned that as trimming had minimal effect on the overall number of SNP calls made, then if it was to be performed, it should at least be done quickly. While no formal assessment of runtime was made in this study, our experience was that there was little reason to use a comparatively slow trimmer, and so fastp, previously benchmarked as up to 5x faster than Trimmomatic and Cutadapt (7) (and with a richer feature set), was greatly preferred.

In terms of user convenience, both TrimGalore and fastp also automatically detect adapter sequence from the reads themselves, whereas Atropos and Trimmomatic require that the adapter be specified, assuming it is known (while Atropos can also predict which adapter sequence is present in a set of reads, this function is included as a separate module; unlike TrimGalore and fastp, Atropos does not perform adapter detection and removal simultaneously). Other fastq pre-processing procedures are also in routine use, such as sequencing error correction (20–23) and the depletion of human contaminants (24), and although the impact these have upon SNP calling is beyond the scope of this study, we cannot exclude the possibility that they confer greater benefit than simply trimming reads. As such, despite the results of the present study, we cannot simply recommend that prior to SNP calling no fastq pre-processing operations be performed at all.

Finally, we need to consider an appropriate set of trimming criteria. It was not our intention to exhaustively test all possible ways to trim a read and so we restricted analysis to 3’ quality-trimming across a range of stringencies, by far the most commonly applied operation (because Illumina reads degrade in quality towards the 3’ end). We found that the greatest apparent benefit to SNP calling performance (evaluated as an increase in F-score) was when using fastp to trim 3’ bases at Phred score thresholds of 20 or lower (as illustrated in **Figure 1**), although would re-iterate that the difference in F-scores when SNP calling using untrimmed data, and data trimmed using these approaches, was statistically insignificant. It is also important to note while the four read trimmers used in this study share the same core functionality of adapter- and quality-trimming, they differ in several respects, having different feature sets and, in the case of fastp, implementing several additional filters by default (which we did not explicitly disable). Irrespective of additional 3’ quality-trimming, by default fastp automatically detects and remove adapters, discards reads with > 5 N bases, performs polyG trimming (should it detect NextSeq or NovaSeq input), and requires a minimum ‘qualified quantity’ per read, i.e. that > 40% of the bases in each read have Phred > 15 (disabling the ‘qualified quantity’ filter renders the performance of fastp similar to other trimmers, especially in terms of precision; see **Supplementary Figure 3**). That more filters are applied by default explains why fastp is the only trimmer for which recall (and thereby F-score) was found to increase after trimming (**Supplementary Figure 2**). By contrast, the other three trimmers, in only performing 3’ trimming, more directly affect an improvement to precision (**Supplementary Figure 1**) although as this occurs at the expense of recall, the overall effect on F-score appears detrimental (note, however, that in absolute terms, these differences were statistically insignificant; **Figure 1**).

For any given trimming program, it is possible to apply the same set of filters, as well as to add others, such as trimming leading as well as trailing bases or to clip reads should the average quality within a sliding window advanced from the 5’ to 3’ end fall below a minimum threshold (a feature of Trimmomatic and fastp, but not TrimGalore or Atropos). However, although drawing from a richer set of filters seems intuitively superior, in practice they are unlikely to make a substantive difference. We have already shown that even after removing > 20% of the (lower-quality) bases from *E. coli*, *M. tuberculosis* and *S. aureus* reads, essentially the same SNPs are still called. It is reasonable to believe that extra filters would for the most part act as complementary methods of removing the same set of lower-quality bases. By way of illustration, we re-calculated F-score for the set of Gram-negative bacteria after pre-processing each sample using fastp with four parameters in addition to those used previously (the default settings, plus 3’ quality-trimming; see **Supplementary Table 2**). Compared to the previous use of fastp (i.e. to perform adapter-trimming, 3’ quality trimming, and to require a minimum read length of 50bp and a ‘qualified quantity’ of bases), the addition of four extra parameters, including base correction and a ‘low complexity’ filter, made no significant difference to SNP calling (the correlation between F-scores when using the two sets of filters was Spearman’s *rho* = 0.999, p < 2.2 × 10^−16^; **Supplementary Figure 6**).

In general, overly conservative filters may prove counterproductive. Especially stringent quality-trimming (for instance, Q < 30) could artificially reduce coverage depth by discarding a larger proportion of reads, as well as shortening many more. In the absence of an additional filter on the basis of minimum read length, reads that are too short are also more likely to be misaligned. This is consistent with a previous study which explored the effects of RNA-seq read trimming upon gene expression estimates, finding that the majority of differences were driven by the spurious mapping of short reads, which could be mitigated by requiring a minimum read length (25). The default minimum read lengths for fastp and TrimGalore are 15 and 20bp, respectively, both relatively short given current Illumina read lengths (> 300bp). In this study, we consistently required a minimum read length of 50bp, one third of the length of the shortest raw read in any sample, 150bp. It would in principle be possible to use an aligner optimised for comparatively short reads, such as Bowtie (26), to perhaps more accurately map those truncated by stringent trimming, although far more pragmatic to simply abstain from such stringency to begin with. As **Figure 1** illustrates, overall SNP calling performance can actually decrease when using overly conservative filters: relative to untrimmed data, F-score is notably reduced when using Atropos and TrimGalore with Q thresholds of 30, a consequence of marginally improving precision (**Supplementary Figure 1**) at the more substantial expense of recall (**Supplementary Figure 2**). Consistent with this, a previous study has advocated a more gentle trimming strategy (Q < 5) as optimal across a range of metrics (6). At least relative to especially stringent trimming, this conclusion appears supported by **Figure 1** – essentially the same distribution of F-scores can be seen for Q < 5 as for Q < 20. However, this is to not to say that, prior to SNP calling, gentle trimming is necessarily superior to no trimming at all, as in our initial Gram-negative dataset the absolute difference in F-score for untrimmed reads relative to reads trimmed using Q < 5 was statistically insignificant (Mann-Whitney U p = 0.98). However, when re-analysing the larger set of publicly-archived *E. coli* reads, we found that the percentage of SNPs identically called before and after trimming was fractionally higher when trimming at Q < 5 compared to Q < 20 (median percentage 99.91% and 99.87%, respectively; Mann-Whitney U p < 2.2×10^−16^; **Supplementary Table 4**). This suggests that more rigorous trimming does allow a small number of additional non-identical calls to be made – although whether these additional calls represent true SNPs not detectable using untrimmed or lightly-trimmed data, or simply additional errors, cannot be determined. Regardless, the percentage of non-homozygous calls (considered a proxy of false positives) was not significantly different between the Q < 5 and Q < 20 data (median percentage 9.02% and 9.05%, respectively; Mann-Whitney U p = 0.96). Many ‘general purpose’ pre-processing tools perform similar functions to those used in this study, including AdapterRemoval v2 (27), AfterQC (28), AlienTrimmer (29), Btrim (30), fastQ_brew (31), FastqPuri (32), ngsShoRT (33), PEAT (34), SeqPurge (35), SeqTrim (36), Skewer (37), and SOAPnuke (38), alongside more protocol-specific tools such as NxTrim (39) and NextClip (40), both of which were designed to remove adaptors from Illumina Nextera mate pairs. We did not seek to evaluate a comprehensive range of tools because our primary concern was not to identify the highest-performing read trimmer *per se*, but to explore the effect of trimming, in general, upon bacterial SNP calling accuracy. We anticipate our findings would be generalizable to a broad range of read trimmers as these essentially share the same core functionality although differ in runtime and memory use.

## Conclusions

A simple means of improving any particular SNP calling pipeline is to remove minimal-value operations, as this decreases the computational time and data manipulation required. The benefit of various operations may not be universally realised and so in many situations could prove an unnecessary computational expense. This has previously been demonstrated for several post-alignment processing steps, such as local indel realignment and base quality score recalibration, when calling variants from exome sequencing data (41). A previous study also demonstrated that PCR duplicate removal had minimal effect on SNP calling and questioned its necessity as a routine procedure (42) (it has not escaped our notice that we duplicate-mask in our own pipelines; this perhaps reflects the ease with which such analytic habits become ingrained).

By re-analysing > 6500 publicly-archived sequencing datasets from *E. coli*, *M. tuberculosis* and *S. aureus*, we found that the retention of lower-quality bases and residual adapter contaminants had minimal effect upon SNP calling. Of the approximately 125 million SNPs called across all samples, 98.8% were identically called irrespective of whether raw reads or trimmed reads were used. This suggests that quality- and adapter-trimming, although routine fastq pre-processing operations, add relatively little value to a SNP calling pipeline and may only be necessary if small differences in the absolute number of SNP calls are critical (such as, for instance, when predicting antimicrobial resistance).

As such, given the majority of read trimmers perform the same basic functions with comparable accuracy, there seems little practical reason to make use of any other than the fastest (currently fastp (7)), if at all. Our results also suggest that if pre-processing is performed then in terms of optimal parameters, we would consider not being too conservative (trimming at a quality threshold of 20 or lower), not retaining reads unpaired as a consequence of trimming, and not relying on 3’ trimming alone (as shown with fastp, there appeared greater performance when supplementing 3’ trimming with a minimum ‘qualified quantity’ per read).

Read trimming remains routinely performed prior to SNP calling likely out of concern that to do otherwise would substantially increase the number of false positive calls. While historically this may have been the case, our data suggests this concern is now unfounded.

## Materials and Methods

### Evaluating the effect of read trimming on SNP calling accuracy

To evaluate the effect of read trimming upon SNP calling, we first required a truth set of SNPs against which comparisons could be made. These were generated as previously described (16) and briefly recapitulated here. Firstly, a dataset was obtained from a previous study (15) that comprised 17 parallel sets of 150 bp Illumina HiSeq 4000 paired-end short reads and both Oxford Nanopore Technologies (ONT) and SMRT Pacific Biosciences (PacBio) long reads for 4 *Enterobacter* spp., 4 *Klebsiella* spp., 4 *Citrobacter* spp., and 3 *Escherichia coli* (all environmentally-sourced), plus subcultures of stocks from 2 reference strains, *K. pneumoniae* subsp. *pneumoniae* MGH 78,578 and *E. coli* CFT073. Sample accessions are listed in **Supplementary Table 1**.

To assess how divergence between the source and reference genomes affected the performance of SNP calling pipelines, we used this Gram-negative dataset to generate truth sets of SNPs. To do so, whole genome alignments were made between each closed assembly and each species’ reference genome using both nucmer (from MUMmer v4.0.0beta2) (43) and Parsnp v1.2 (44) with default parameters, with common SNPs then identified within one-to-one alignment blocks. The set of SNPs identically called by both nucmer and Parsnp in both *de novo* assemblies were considered the ‘truth set’ and used to evaluate the performance of each trimmer/aligner/caller pipeline, as follows.

We first adapter- and quality-trimmed each of the 17 sets of Illumina reads using four different read trimmers – Atropos v1.1.25 (9), fastp v0.20.1 (7), TrimGalore v0.5.0 (https://github.com/FelixKrueger/TrimGalore, accessed 1^st^ April 2020) and Trimmomatic v0.38 (8) – as well as retaining an ‘untrimmed’ control. Read trimmers often have rich feature sets although our concern with this study was not to systematically evaluate the broad range of parameters by which a read may be trimmed but to assess, in general, the added-value benefit of trimming when a minimum-effort application was made for each program. In this respect, we considered ‘trimming’ to encompass two simultaneous pre-processing operations: the removal of adapter sequence (using default parameters where possible) and 3’ quality-trimming (across a range of stringencies).

We configured each trimmer to automatically detect and remove adapter sequence, where this was the default setting (fastp, TrimGalore), or to remove the Illumina universal adapter, should specification be required (Atropos, Trimmomatic). As Illumina reads degrade in quality towards the 3’ end, 3’ trimming is by far the most commonly applied pre-processing operation. Alongside adapter removal, we removed trailing bases should they fall below Phred quality thresholds of 2, 5, 10, 20 or 30 (the Phred scale being logarithmic, this represents base call accuracies of 37%, 68%, 90%, 99%, and 99.9%, respectively). We also required a minimum post-trimming read length of 50bp and, where possible, for each trimmer to output both paired and unpaired reads, i.e. those reads where both ends of a pair are retained after trimming, and those where one end is discarded (by default, fastp and TrimGalore discard the entire pair if only one end meets the acceptance criteria). Finally, we did not explicitly disable any filter criterion implemented automatically, considering this to be the default recommendation for general-purpose use. The specific parameters used for each trimmer are detailed in **Supplementary Table 2**.

Using the trimmed (or, as a control, untrimmed) Illumina reads, and the reference genome for each species (listed in **Supplementary Table 1**), we then called SNPs using the pairwise combination of the aligner BWA-mem v0.7.17 (45) with three variant callers, LoFreq v2.1.2 (12), mpileup v1.7 (13), and Strelka v2.9.2 (14), each used with default parameters. Each pipeline applied a common set of post-processing steps: BAM files were cleaned, sorted, had duplicate reads marked, and were indexed using Picard Tools v2.17.11 (46), and VCFs were regularised using the vcfallelicprimitives module of vcflib v1.0.0-rc2 (https://github.com/ekg/vcflib, accessed 1^st^ April 2020). Finally, VCFs were filtered using BCFtools v1.7 (47) to retain only biallelic SNPs with call quality > 20 and > 5 reads mapped at that position, >75% of which, including at least one in each direction, supporting the alternative allele (as in (48), and broadly similar to those recommended by a previous study for maximising SNP calling precision (19)).

To evaluate each pipeline, we calculated precision (positive predictive value), recall (sensitivity) and F-score, a summary measure which considers precision and recall with equal weight, producing a value between 0 and 1 (perfect precision and recall). Precision was calculated as TP/(TP+FP), recall as TP/(TP+FN), and F-score as 2 × ((precision × recall) / (precision + recall)), where TP, FP and FN are the number of true positive, false positive, and false negative SNP calls, respectively.

The command lines used for these pipelines were also previously implemented (16) within a suite of Perl scripts (so as to handle subsidiary data manipulation operations and calculate summary statistics), available at https://github.com/oxfordmmm/GenomicDiversityPaper. The SNP call truth sets are available via the GigaDB repository at http://dx.doi.org/10.5524/100694.

Finally, when SNP calling using publicly-sourced *E. coli*, *M. tuberculosis* and *S. aureus* data (see below), we used only one of the above pipelines, fastp (with trailing Q < 20) / BWA-mem / mpileup, aligning all reads relative to *E. coli* K-12 substr. MG1655 (RefSeq assembly accession GCF_000005845.2), *M. tuberculosis* H37Rv (RefSeq assembly accession GCF_000195955.2) and *S. aureus* subsp. *aureus* NCTC8325 (RefSeq assembly accession GCF_000013425.1), respectively. This fastp / BWA-mem / mpileup pipeline was as previously used upon the Gram-negative dataset, although to reduce runtime was modified to omit the BAM cleaning (i.e. Picard CleanSam), VCF regularisation, and VCF filtering steps (regularisation in any case only necessary when comparing VCFs produced by different variant callers). Command lines are detailed in **Supplementary Table 3**.

### Evaluating the effect of trimming upon a diverse range of publicly-archived sequencing datasets

To obtain a broad range of sequencing data across multiple laboratories, we downloaded the daily-updated SRA BioProject summary file (n = 417,689 BioProjects; ftp://ftp.ncbi.nlm.nih.gov/bioproject/summary.txt, accessed 22^nd^ March 2020), parsing it to extract BioProject IDs with a data type of ‘genome sequencing’ and associated NCBI taxonomy IDs of 562 (*Escherichia coli*), 1773 (*Mycobacterium tuberculosis*) and 1280 (*Staphylococcus aureus*). These are ‘top level’ taxonomy ID, encompassing samples for which an additional level of strain-specificity could not be made. These ‘top level’ IDs were chosen so that, alongside the criteria detailed below, a set of samples would be obtained that was large enough to draw sound conclusions (*E. coli*, *M. tuberculosis* and *S. aureus* being three of the more commonly sequenced bacteria) but not so large as to be computationally expensive to analyse (which would be the case if obtaining all WGS data from every *E. coli*, *M. tuberculosis* and *S. aureus* strain). These three species were chosen to represent different degrees of genomic diversity, from, broadly speaking, high (*E. coli*) to low (*M. tuberculosis*). We used the Entrez Direct suite of utilities (https://www.ncbi.nlm.nih.gov/books/NBK179288/, accessed 1st May 2019) to associate each BioProject ID with a list of SRA sample and run IDs (a ‘RunInfo’ file). RunInfo files were parsed to retain only those runs where ‘Platform’ was ‘ILLUMINA’, ‘Model’ (i.e. sequencer) contained ‘HiSeq’ or ‘MiSeq’ (all of which use TruSeq3 adapters), ‘LibrarySource’ was ‘GENOMIC’, ‘LibraryStrategy’ was ‘WGS’, ‘LibraryLayout’ was ‘PAIRED’, ‘LibrarySelection’ was ‘RANDOM’, ‘avgLength’ was ≥ 150 (i.e., mean read length of 150bp) and ‘spots’ was > 1 and < 5 (i.e., approximating a read depth of > 1 and < 5 million reads). The upper limit on read depth was chosen to minimise the computational cost of processing, although read depth remains sufficiently high as to not compromise SNP calling. For instance, as the *E. coli* genome is approx. 5 million bp in length, sequencing 1-5 million 150bp paired-end reads represents a mean base-level coverage of approx. 60- to 300- fold. These criteria generated sets of 1661 *E. coli*, 4416 *M. tuberculosis* and 1156 *S. aureus* sequencing read sets, detailed in **Supplementary Tables 3, 5 and 7**, respectively. Publicly archived sequencing data often does not contain computationally-accessible metadata regarding any informatic pre-processing and nor is it immediately apparent whether pre-processing has already been performed. As such, we excluded from consideration all samples where < 1% of the original bases could be trimmed, reasoning that in these circumstances the sample had already been pre-processed using broadly similar trimming criteria to those applied here. The final *E. coli* dataset comprised 1606 samples, representing a particularly diverse set of strains. Relative to the *E. coli* reference, K-12 substr. MG1655, the majority of samples contained between approximately 10,000 and 100,000 SNPs, although with outliers containing as few as 15 SNPs and as many as 308,000 (> 103 million SNPs in total), with a mean of 64,476 SNPs per sample (**Supplementary Table 4**). The final *M. tuberculosis* dataset comprised 3946 samples, incorporating subsets from two large-scale studies from South Africa (representing > 2000 of the original 4416 samples) (49) and Canada (> 1000 samples) (50). Each sample contained between approximately 170 and 2500 SNPs (c. 5 million SNPs in total). Across all samples, the mean number of SNPs relative to the *M. tuberculosis* reference, H37Rv, was 1238 (**Supplementary Table 6**). The final *S. aureus* dataset comprised 1100 samples, each of which contained between approximately 1000 and 60,000 SNPs relative to the *S. aureus* reference genome, NCTC8325 (mean 15,167 SNPs per sample, with c. 17 million SNPs in total) (**Supplementary Table 8**).

## Supplementary Figures

**Supplementary Figure 1.**
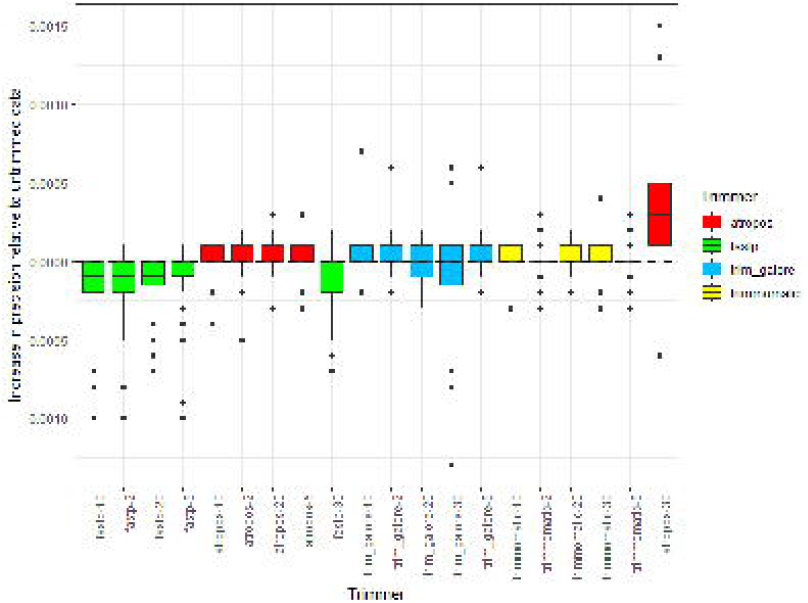
Effect of read trimming upon precision (positive predictive value) when SNP calling in a curated Gram-negative dataset. Median difference in precision per trimmer relative to untrimmed data, across a range of trimming stringencies (i.e. varying the Phred score threshold for trimming 3’ bases). Boxes represent the interquartile range of precision, with midlines representing the median. Upper and lower whiskers extend, respectively, to the largest and smallest values no further than 1.5x the interquartile range. Data beyond the ends of each whisker are outliers and plotted individually. Columns are ordered according to median precision and coloured according to the trimmer used. The dashed line y=0 is marked in black. The raw data for this figure is available in **Supplementary Table 2**. Note that fastp implements quality filters other than 3’ trimming by default, which for the data in this figure were retained. A version of this figure with these filters disabled is available in **Supplementary Figure 3**.

**Supplementary Figure 2.**
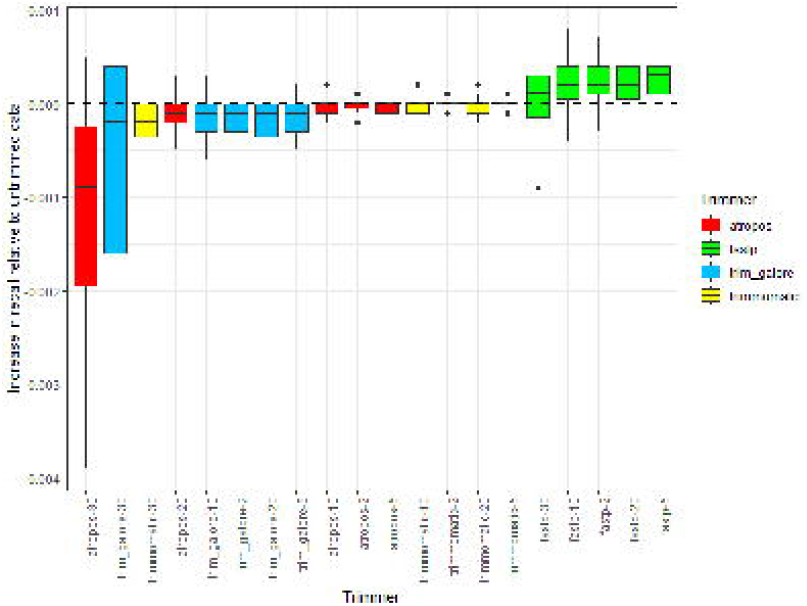
Effect of read trimming upon recall (sensitivity) when SNP calling in a curated Gram-negative dataset. Median difference in recall per trimmer relative to untrimmed data, across a range of trimming stringencies (i.e. varying the Phred score threshold for trimming 3’ bases). Boxes represent the interquartile range of recall, with midlines representing the median. Upper and lower whiskers extend, respectively, to the largest and smallest values no further than 1.5x the interquartile range. Data beyond the ends of each whisker are outliers and plotted individually. Columns are ordered according to median recall and coloured according to the trimmer used. The dashed line y=0 is marked in black. The raw data for this figure is available in **Supplementary Table 2**. Note that fastp implements quality filters other than 3’ trimming by default, which for the data in this figure were retained. A version of this figure with these filters disabled is available in **Supplementary Figure 3**.

**Supplementary Figure 3.**
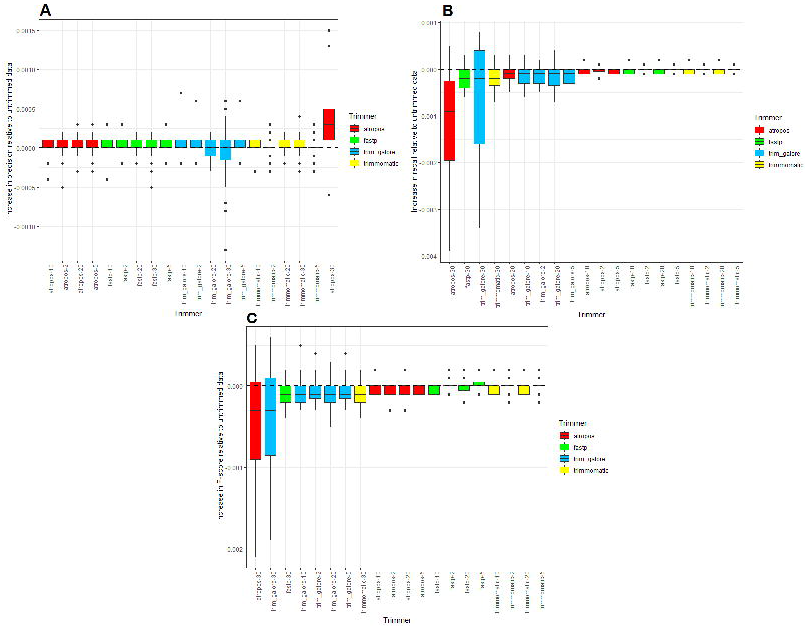
Effect of read trimming upon precision (positive predictive value), recall (sensitivity) and F-score when SNP calling in a curated Gram-negative dataset, disabling fastp default options. The data shown in this figure is as shown in **Figure 1** (F-score), **Supplementary Figure 1** (precision) and **Supplementary Figure 2** (recall), with the exception of values for fastp, in which the default trimming option (‘qualified quantity’, requiring that > 40% of the bases in each read have Phred > 15) is now explicitly disabled and the trimming performed is limited only to adapter removal and 3’ quality trimming. This figure shows that across all four trimmers, 3’ quality-trimming, in general, marginally increases precision (panel A) but at the expense of recall (panel B) and thereby overall F-score (panel C). Panels show the median difference in precision (A), recall (B) or F-score (C) per trimmer relative to untrimmed data, across a range of trimming stringencies (i.e. varying the Phred score threshold for trimming 3’ bases). Boxes represent the interquartile range of precision (A), recall (B) and F-score (C), with midlines representing the median. Upper and lower whiskers extend, respectively, to the largest and smallest values no further than 1.5x the interquartile range. Data beyond the ends of each whisker are outliers and plotted individually. Columns are ordered according to median precision (A), recall (B) and F-score (C), and coloured according to the trimmer used. The dashed line y=0 is marked in black. The raw data for this figure is available in **Supplementary Table 2**.

**Supplementary Figure 4.**
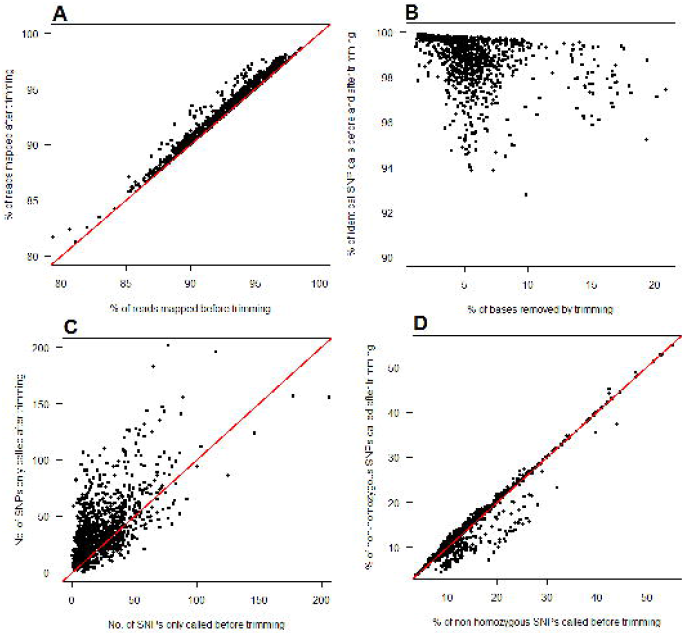
Effect of read trimming upon SNP calls made using publicly-archived *S. aureus* sequencing data. This figure recapitulates patterns seen in **Figure 2** and illustrates the effect of read trimming upon SNP calls made in *S. aureus*. Trimming marginally increases the proportion of successfully aligned reads, albeit from a high baseline value, > 85% (panel A). The majority of SNPs (> 96%) are nevertheless called irrespective of any trimming (panel B). A relatively small number of SNPs (often < 200) are only called when using either raw, or trimmed data, but not both (panel C). The proportion of non-homozygous SNP calls, considered a proxy of false positive calling, decreases when using trimmed data (panel D). The raw data for this figure is available in **Supplementary Table 8** and represents 1100 *M. tuberculosis* samples, with a mean of 15,167 SNPs per sample. The red line denotes y = x.

**Supplementary Figure 5.**
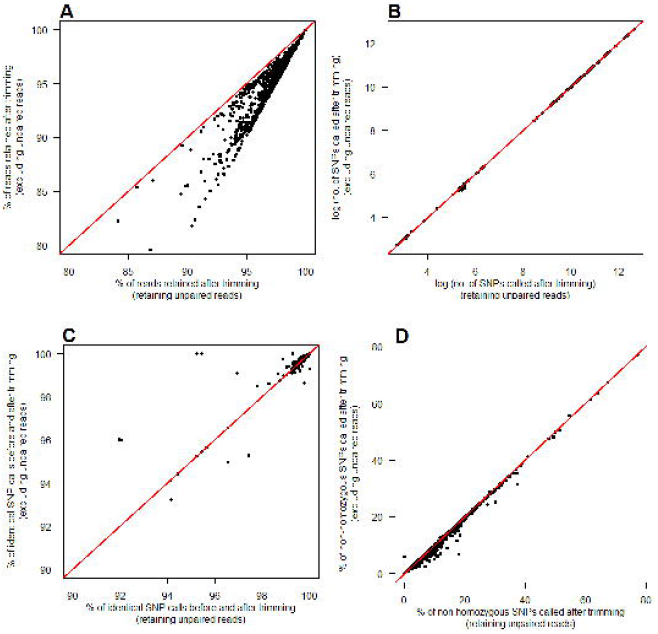
Discarding unpaired reads after trimming has negligible impact on the number of SNPs called in *E. coli*. This figure illustrates the effect of read trimming upon SNP calls made in *E. coli*, with the read trimmer retaining or discarding unpaired reads (unpaired reads are those when one end of a pair is discarded by the trimmer, and so are output as *de facto* single-end; SE reads). While discarding SE reads reduces, by definition, the proportion of reads available for mapping and SNP calling, this only represents a substantial proportion (> 5%) of the total in limited cases (panel A). There is little discernible difference both in the absolute number of SNPs called with and without SE reads (panel B) and the percentage of SNPs identically called before and after trimming (panel C), although seemingly a small decrease in the number of non-homozygous calls (a proxy of false positives) made when discarding the unpaired reads (panel D). Raw data for this figure is available in **Supplementary Table 4** and represents 1606 *E. coli* samples. The red line denotes y = x.

**Supplementary Figure 6.**
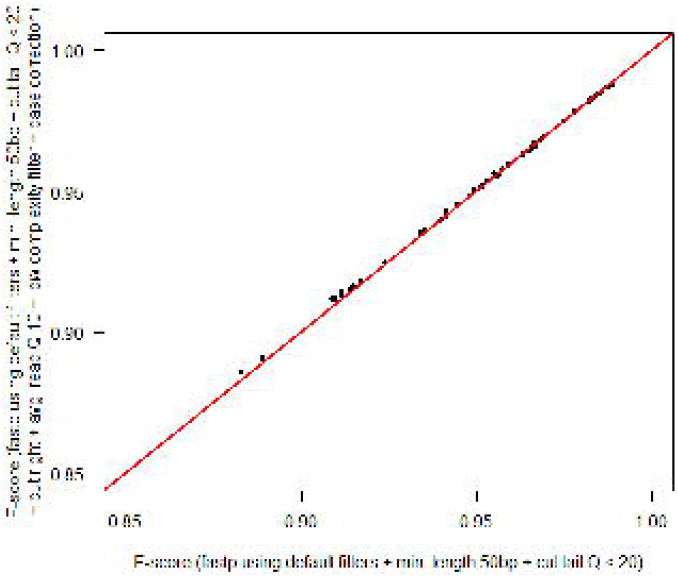
Pre-processing with additional filters has negligible effect on F-score when SNP calling in a curated Gram-negative dataset. This figure shows the F-score per aligner/caller combination when SNP calling each of 17 Gram-negative genomes, after pre-processing reads using fastp with two sets of parameters: one simple, one complex. The x-axis applies a simple set of filters: the two default settings of adapter-trimming and a minimum ‘qualified quantity’ of bases, plus 3’ quality trimming (‘cut tail’) and a minimum read length of 50bp. The y-axis supplements these filters with four more: ‘cut left’ (cutting reads should the mean quality within a 4bp window, advanced 5’ to 3’, fall below 20), ‘low complexity’ (requiring that 30% or more of the bases in each read are followed by a different base), ‘average quality’ (requiring a mean base quality across the entire read of > 10), and ‘correction’ (correcting mismatched base pairs in regions of paired end reads that overlap each other, if one base has a quality higher than the other [by default, requiring a minimum overlap of 30bp, a difference in base qualities of 5, and no more than 20% of the bases in the overlapping region needing correction]). The raw data for this figure is available in **Supplementary Table 2**.

## Supplementary Tables

**Supplementary Table 1.** Environmentally-sourced/reference Gram-negative isolates and associated representative genomes.

**Supplementary Table 2.** Performance of trimmer/aligner/caller pipelines across a range of Gram-negative genomes and trimming strategies.

**Supplementary Table 3.** Sources of *E. coli* sequencing data (Illumina paired-end reads of ≥

150bp).

**Supplementary Table 4.** Proportion of SNPs called before and after pre-processing 1606 publicly-archived *E. coli* sequencing reads.

**Supplementary Table 5.** Sources of *M. tuberculosis* sequencing data (Illumina paired-end reads of ≥ 150bp).

**Supplementary Table 6.** Proportion of SNPs called before and after pre-processing 3946 publicly-archived *M. tuberculosis* sequencing reads.

**Supplementary Table 7.** Sources of *S. aureus* sequencing data (Illumina paired-end reads of

≥ 150bp).

**Supplementary Table 8.** Proportion of SNPs called before and after pre-processing 1100 publicly-archived *S. aureus* sequencing reads.

## Data bibliography

1. Short- and long read sequencing data and assemblies: Illumina, ONT and PacBio sequencing data, as described in a previous study (15), are available via the NCBI Sequence Read Archive under BioProject PRJNA42251 (https://www.ncbi.nlm.nih.gov/bioproject/PRJNA422511) with the associated Illumina/ONT and Illumina/PacBio assemblies available via FigShare (https://doi.org/10.6084/m9.figshare.7649051).

2. SNP call truth sets: the aforementioned Illumina reads were used in a previous evaluation of 209 SNP calling pipelines (16), for which the SNP call truth sets, also used here, are available via the GigaDB repository (http://dx.doi.org/10.5524/100694).

## Authors’ contributions

SJB conceived of and designed the study, performed all analyses, and wrote the manuscript.

